# Robust and generalizable segmentation of human functional tissue units

**DOI:** 10.1101/2021.11.09.467810

**Authors:** Leah L. Godwin, Yingnan Ju, Naveksha Sood, Yashvardhan Jain, Ellen M. Quardokus, Andreas Bueckle, Teri Longacre, Aaron Horning, Yiing Lin, Edward D. Esplin, John W. Hickey, Michael P. Snyder, N. Heath Patterson, Jeffrey M. Spraggins, Katy Börner

**Author notes:** **Corresponding authors**, Leah L. Godwin, Katy Börner.

## Abstract

The Human BioMolecular Atlas Program aims to compile a reference atlas for the healthy human adult body at the cellular level. Functional tissue units (FTU, e.g., renal glomeruli and colonic crypts) are of pathobiological significance and relevant for modeling and understanding disease progression. Yet, annotation of FTUs is time consuming and expensive when done manually and existing algorithms achieve low accuracy and do not generalize well. This paper compares the five winning algorithms from the “Hacking the Kidney” Kaggle competition to which more than a thousand teams from sixty countries contributed. We compare the accuracy and performance of the algorithms on a large-scale renal glomerulus Periodic acid-Schiff stain dataset and their generalizability to a colonic crypts hematoxylin and eosin stain dataset. Results help to characterize how the number of FTUs per unit area differs in relationship to their position in kidney and colon with respect to age, sex, body mass index (BMI), and other clinical data and are relevant for advancing pathology, anatomy, and surgery.

## Introduction

The Human BioMolecular Atlas Program (HuBMAP) aims to create an open, computable human reference atlas (HRA) at the cellular level^1^. The envisioned HRA will make it possible to register and explore human tissue data across scales—from the whole-body macro-anatomy level to the single-cell level. Medically and pathologically relevant functional tissue units (FTUs) are seen as important for bridging the meter level scale of the whole body to the micrometer scale of single cells. Functional tissue units are defined as a three-dimensional block of cells centered around a capillary where each cell is within diffusion distance from any other cell within the same block; a term coined by De Bono et al.^2^ Plus, FTUs accomplish important biomedical functions and are “units of physiological function that are replicated multiple times in a whole organ^3^”. The value of FTUs is acknowledged by the scientific and medical communities, yet limited data exists about human diversity in terms of the number and size distribution for a single organ and across individuals with different age, sex, BMI. A key reason for this knowledge gap is the fact that annotation of FTUs is time consuming and expensive when done manually. For example, there are over 1 million glomeruli in an average human kidney^3^, but a trained pathologist needs ca. 10 hours of time to annotate 200 FTUs. FTU detection algorithms exist^4–12^ and approaches range from simple thresholding^11^ to deep learning methods. Existing methods achieved varying levels of performance (see **Supplementary Tables 1 and 2** and performance metric definitions in Methods) and face challenges when applied to human data (e.g., training on murine glomerulus data generated false positives when applied to the much larger glomeruli in human data^12^). Rapid progress is desirable as a robust and highly performant FTU detector would make it possible to compute size, shape, variability in number and location of FTUs within tissue samples and to use this information to characterize human diversity—providing critical information for the construction of a spatially accurate and semantically explicit model of the human body.

This paper is organized as follows: We present results from comparing the top-five winning algorithms from the recent “Hacking the Kidney” Kaggle competition^13^. Specifically, we reproduce results and then apply the five algorithms to segment colon data (from scratch and transfer learning) to determine their generalizability to other FTU types. Segmentation data is then used to characterize the number of FTUs per unit area in dependence on location in the human body as well as donor sex, age, and BMI. Last but not least, we discuss how FTU detection advances the construction of a Human Reference Atlas. All data and code can be freely accessed at https://github.com/cns-iu/ccf-research-kaggle-2021.

## Results

### Data Preparation

For the “Hacking the Kidney” Kaggle competition, a unique dataset was compiled comprising 30 Periodic acid-Schiff (PAS) stain whole slide images (WSI) with 7,102 annotated renal glomeruli (see **Supplementary Table 3** and Methods). To determine if algorithms generalize to other FTU types, a second dataset was compiled comprising seven colon hematoxylin and eosin stain WSI with 395 segmented colonic crypts (see **Supplementary Table 4** and Methods). **Fig. 1a** shows the tissue extraction sites for the 30 kidney and seven colon WSI datasets (explore three-dimensional reference organs at https://cns-iu.github.io/ccf-research-kaggle-2021). Exemplary glom and crypt annotations are given in **Fig. 1b. Fig. 1c** lists basic information (sex, age, BMI) for all 37 datasets; the datasets are sorted by their spatial location in the reference organ— using the mass point of the tissue block from which they were extracted (top-most block is on top), see Methods.

**Figure 1.**
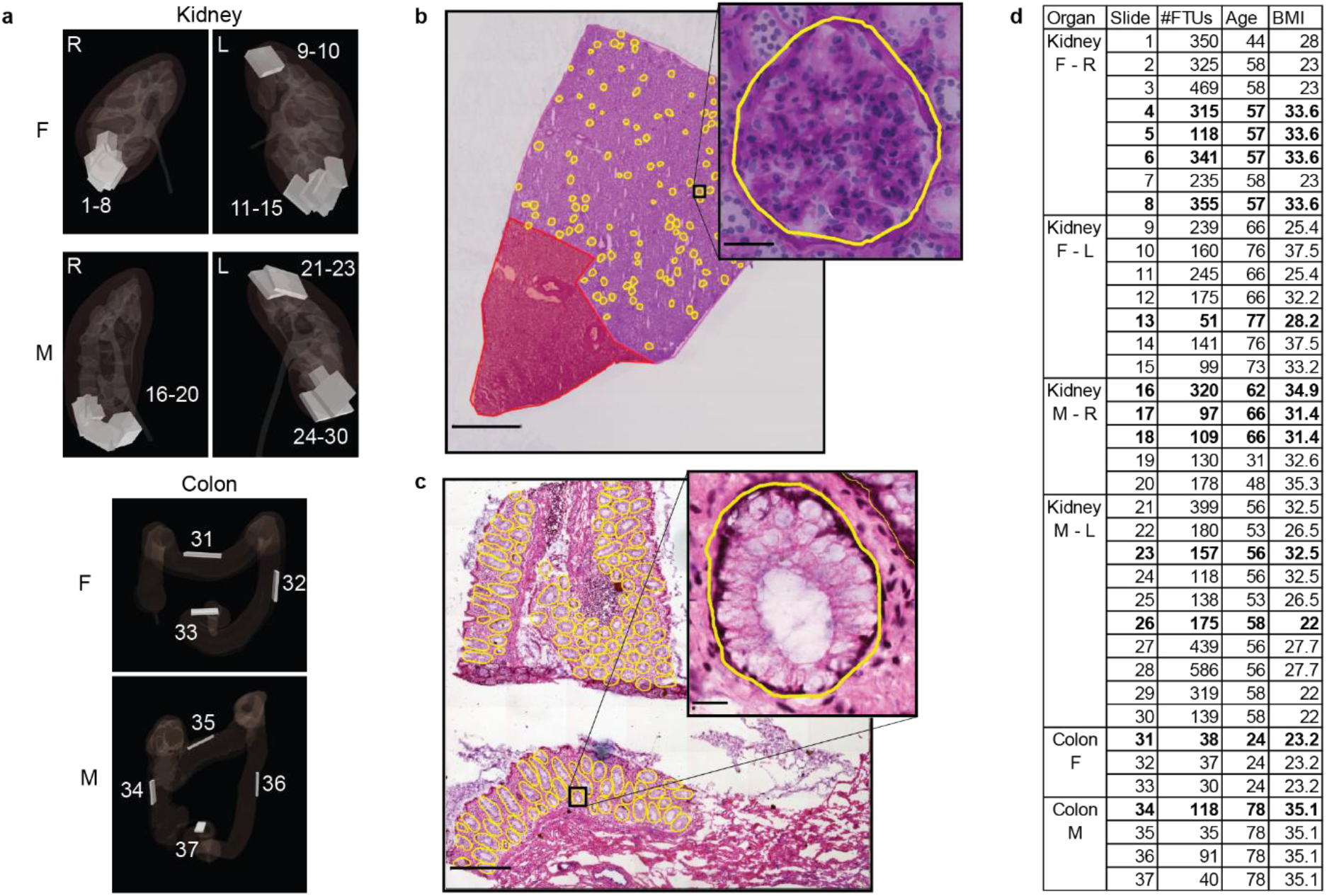
FTU Datasets. **a.** The 30 kidney and 7 colon tissue datasets were registered into the corresponding male/female, left/right HuBMAP 3D reference organs for kidney and colon to capture the size, position, and rotation of tissue blocks. **b.** Sample kidney WSI (scale bar: 2mm) with zoom into one glomerulus annotation (scale bar: 50μm). Right below is a sample colon WSI (scale bar: 500μm) with zoom into a single crypt annotation (scale bar: 20μm). **c.** Metadata for 37 WSI sorted top-down by vertical location within the reference organs; test datasets are given in bold.

The Kaggle dataset was split into a 15 WSI training and 5 WSI validation dataset; both were available to competition participants. The 10 WSI private test dataset was used for scoring algorithm performance, see competition design in Methods. Analogously, the colon dataset was split into five WSI used in training and two WSI used for testing. All test datasets are rendered in bold in **Fig. 1c**.

### Algorithm Comparison

The top-5 winning algorithms from the “Hacking the Kidney” Kaggle Competition are from teams named Tom, Gleb, Whats goin on, DeepLive.exe, and Deepflash2. All five use the UNet architecture, see algorithm descriptions in Methods section. Performance results are shown in **Fig. 2** using violin plots for three metrics: DICE, precision, and recall (see details in the Methods section, data values are in **Supplementary Table 5** and interactive data visualization at https://cns-iu.github.io/ccf-research-kaggle-2021). For each of the five algorithms we report DICE coefficient in **Fig. 2a**, recall in **Fig. 2b**, and precision in **Fig. 2c**. For each metric, we show distribution for the ten kidney WSI with 2038 glomeruli on the left and the distribution for the two colon WSI with 160 crypts transfer learning predictions on the right. Performance on kidney vs. colon data can be easily compared. As expected, all five algorithms have a higher DICE coefficient for kidney data than for transfer learning on colon data. Tom—the Kaggle competition performance winner—has the highest mean DICE score of 0.88 for transfer learning on colon data. As for recall, Tom again has the highest value with 0.92—with 9 false negatives and 17 false positives out of 160 crypts. In terms of precision, DeepLive.exe wins with 0.86—with 8 false negatives and 19 false positives. The data in **Supplementary Table 5** also shows that all five algorithms have the lowest DICE scores on WSI 7 and 28. 7 has a low number of crypts, only 51; any false positive/negative prediction has a major impact on the DICE coefficient. WSI 28 has several artifacts and overall lower quality (higher saturation and darker) than other kidney WSIs. The crypt segmentation solution for Tom in comparison with ground truth for colon data can be explored at https://cns-iu.github.io/ccf-research-kaggle-2021.

**Figure 2.**
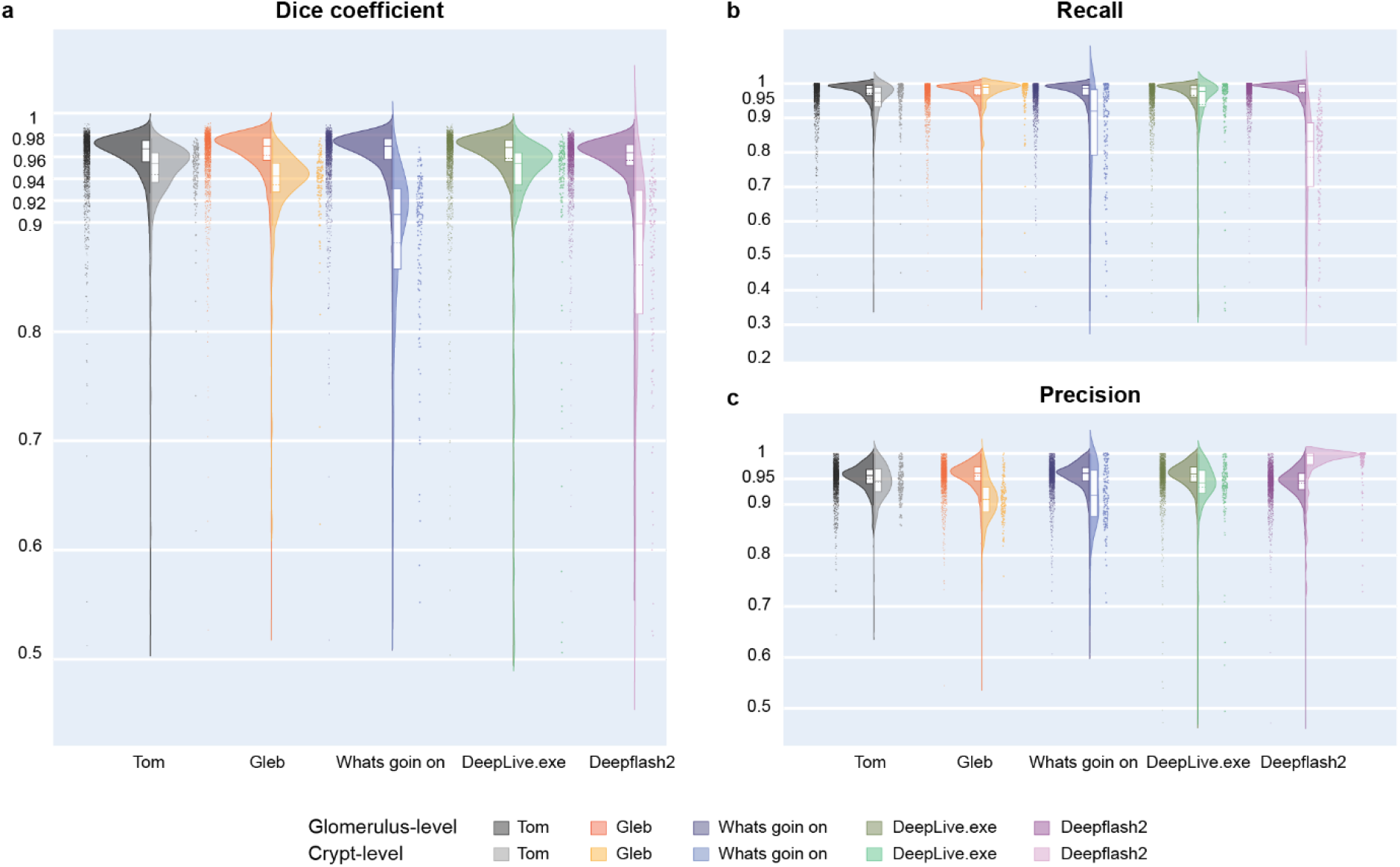
Algorithm Performance Results. Violin plots show performance for kidney on the left (one dot per 2,038 glomeruli) and transfer learning performance for colon data (one dot for each of the 160 crypts) on the right. **a.** DICE coefficient. **b.** Recall performance. **c.** Precision performance. Interactive versions of these graphs are at https://cns-iu.github.io/ccf-research-kaggle-2021.

Run time performance was recorded for the training phase on kidney data, colon data exclusively (no transfer), and on kidney data and colon data, see **Table 1**. We also report run time for the two prediction tasks: from scratch without transfer learning (i.e., trained on five colon, tested on two colon datasets) and transfer learning (i.e., trained on 15 kidney datasets initially and then trained on five colon datasets, then tested on two colon datasets), see Methods section for details.

**Table 1.** Approximate run time performance for training and prediction runs for all five algorithms. (*Times reported by teams.)

| Model | Training*<br>on Kidney | Train on<br>Colon (No<br>transfer<br>learning) | Train on<br>Colon<br>(transfer<br>learning) | Inference on<br>Kidney Data<br>(n=10) | Inference on<br>Colon Data<br>(n=2) |
| --- | --- | --- | --- | --- | --- |
| Tom | 12 hours | 6 hours | 4 hours | 3 hours | 2 mins |
| Gleb | 8 hours | 4 hours | 4 hours | 3 hours | 2 mins |
| Whats goin on | 26 hours | 3 hours | 3 hours | 30 mins | 2 mins |
| DeepLive.exe | 3 hours | 48 hours | 48 hours | 3 hours | 2 mins |
| Deepflash2 | 5 hours | 50 minutes | 1 hour | 30 mins | 2 mins |

As can be seen, training takes time (three hours to 48 hours) while prediction runs are fast (3-30h for kidney and 2mins for colon). Total algorithm run time (training on kidney, then colon plus inference on colon) is lowest for Deepflash2 (6h), followed by Gleb (12h) and Tom (16h). Whats goin on and Deepflash2 are fast in kidney prediction (30mins).
Note that one of the winning teams (Deepflash2) had access to a biomedical expert as a teammate and two of the teams (Tom and DeepLive.exe) used additional data to improve generalizability. The teams did employ clever approaches to sampling (e.g., Deepflash2 using probabilistic sampling to make training time faster) and classification (e.g., DeepLive.exe using a classifier to distinguish between healthy and diseased glomeruli).

### Characterizing Human Diversity

Information on the spatial location of FTUs in human tissue makes it possible to characterize human diversity in support of understanding human diversity. Specifically, we use data on 7,102 glomeruli and 395 crypt annotations to study the impact of sex, age, BMI but also location of tissue in the human body on the number of FTUs per square millimeter. **Fig. 3a** shows the impact of age on the number of detected glomeruli Blocks that have the same age are from the very same donor. As can be seen, out of the 8 females, one has 4 tissue blocks (in y-sequence, top-down: 5, 6, 8, 4), one has 3 tissue blocks (3, 2, 7), and two have 2 tissue blocks (9, 11; 10,14). For the 8 males, two have 3 tissue blocks (in y-sequence: 21, 23, 24; 29, 30, 26) and three have 2 tissue blocks (25, 22; 28, 27; 18, 17). In general, the number of glomeruli per mm^2^ seems to decrease for females and increase for males (except for HBM 322:KQBK.747) going from top to bottom of the kidney. (Slides are numbered by y-position of the 3D reference organ registration.)

**Figure 3.**
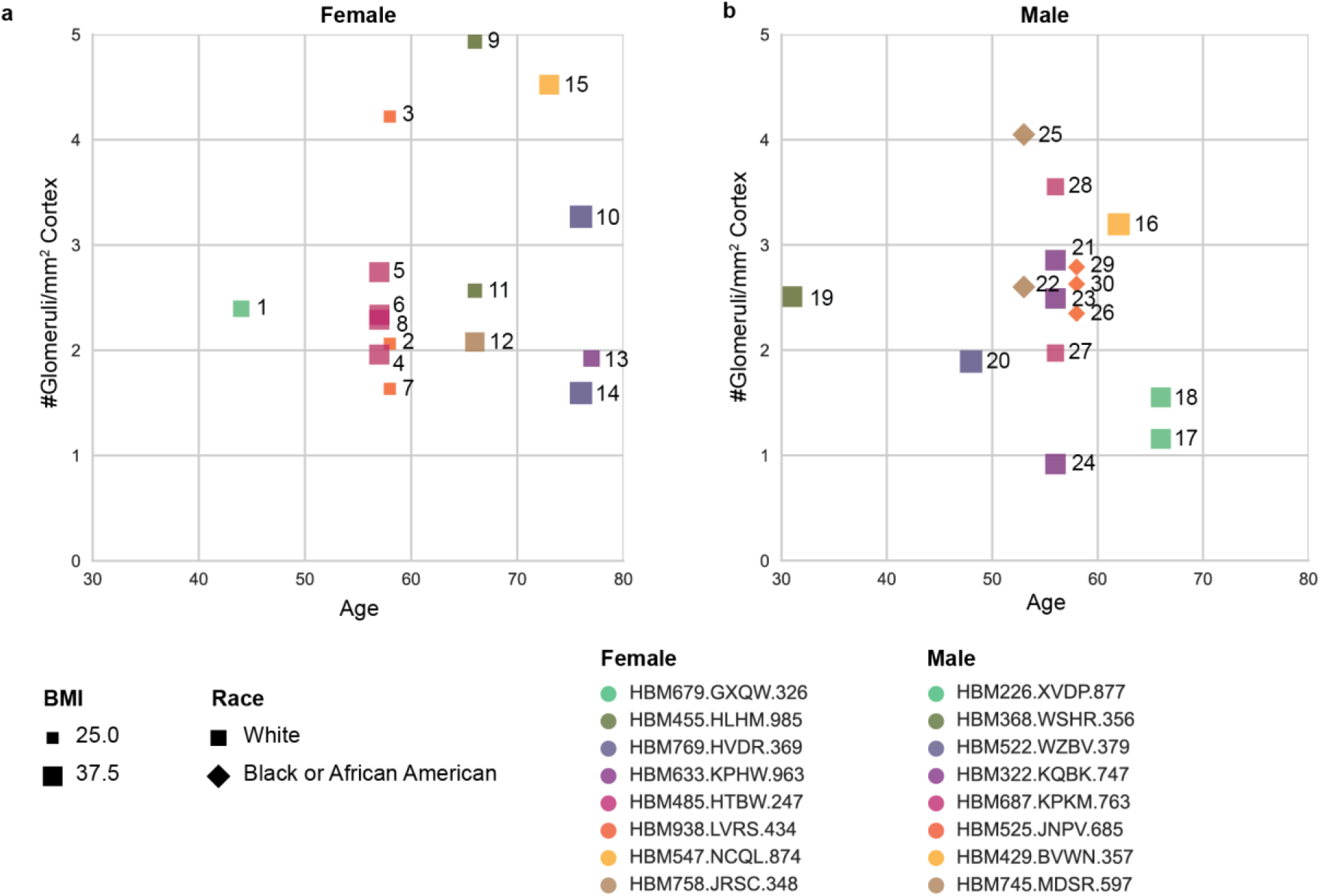
Number of FTUs in dependence of donor sex, age, ethnicity, BMI, and spatial tissue location. The plots show the number of FTUs per square millimeter. Donors are color coded, BMI corresponds to size coding of symbols, squares denote ethnicity with squares indicating White and diamond black or African American, age is position on x-axis. **a.** Graph for kidney, male. **b.** Graph for kidney, female.

Understanding the spatial location and density of FTUs across organs is critically important for advancing the construction of a Human Reference Atlas (HRA)^14^. A robust and highly performant FTU detector would make it possible to compute the size, shape, variability in number, and location of FTUs within tissue samples. This information can then be used to characterize human diversity; to decide on what tissue data should be collected next to improve the coverage and quality of a HRA, and for quality control (e.g., FTU size and density that is vastly different from normal might indicate disease, problems with data preprocess, or segmentation algorithms).

## Discussion

There is a need for efficient and accurate segmentation of FTUs both within HuBMAP and the broader biomedical community. Despite many breakthroughs in the field, the currently available methods for glomerulus and crypt image segmentation do not meet this need. This paper compared winning algorithms from the recently completed “HuBMAP - Hacking the Kidney” Kaggle competition and identified the Tom algorithm as the most accurate, generalizable, and best run time performant algorithm. To our knowledge, this is the first time that scientific evidence is provided of the value of Kaggle competitions to develop algorithms that are superior to existing code. The 1,600 Kaggle teams performed many iterations of experimentation that our team would not have had the time/resources for or thought to try; they build on solutions taken from many different domains to arrive at the winning entries. Given the success of this first competition; we are planning three new Kaggle competitions that aim to advance tissue segmentation and annotation.

Code has been documented and made available freely for anyone to use. We are in the process of preparing this winning algorithm for production usage in the HuBMAP Data Portal^15^ and making it available as part of the HRA ecosystem—for free usage by anyone interested to register and analyze tissue. Going forward, kidney and colon datasets that were spatially registered using the HuBMAP registration user interface^16^ and that have anatomical structures in which FTUs are known to exist will automatically be segmented. In addition, we are in the process of creating additional datasets with FTU annotations for other organs (nephron tubule in kidney; alveoli in lung; hair follicle in skin; white pulp in spleen; lobule in liver; lobule in lymph node; lobule in thymus; sarcomere in heart). The datasets will be used to run transfer learning for FTUs in other organs and to develop robust pipelines for the automatic segmentation and analysis of FTUs across major organs of the human body.

Since all the five winning models use some specific methodology—either in data preprocessing, sampling, or training—that gives them an edge over the others, we are exploring taking the best parts of each and constructing a sixth model. For example, Deepflash2 uses a probabilistic sampling strategy that makes its training faster; DeepLive.exe uses additional data and a classifier in its model to improve its results. Plus, training time can be reduced by using distributed training; training can be monitored in support of optimization and explainability.

Going forward, 3D data of FTUs will be used to identify the number, size, and shape of FTUs in support of machine learning and single-cell simulation of the structure and function of FTUs. Resulting data will be used to increase our collective understanding of (and variability in) the size, number, and location of FTUs in relation to donor sex, age, ethniticy, and BMI. This data and work is also critical for integrating top-down (segmenting out larger known structures) and bottom-up (single-cell data) in multiplexed imaging techniques and relating composition within these structures. Top-down and bottom-up data integration and analysis are needed for constructing an accurate and comprehensive Human Reference Atlas.

## Methods

### Datasets

#### Renal glomeruli data

Renal glomeruli are groups of capillaries that facilitate filtration of blood in the outer layer of kidney tissue known as the cortex^17^. The size of normal glomeruli in humans ranges from 100-350 μm in diameter and they have a roughly spherical shape^4^. Glomeruli contain four cell types: parietal epithelial cells (CL:1000452), podocytes (CL:0000653), fenestrated endothelial cells (a.k.a. glomerular capillary endothelial cell CL:1001005), and mesangial cells (CL:1000742)^18^. Parietal epithelial cells form the Bowman’s capsule. Podocytes cover the outer layer of the filtration barrier. Fenestrated endothelial cells are in direct contact with blood and coated with a glycolipid and glycoprotein matrix called glycocalyx. Mesangial cells occupy the space between the capillary blood vessel loops and are stained by the colorimetric histological stain called Periodic acid-Schiff (PAS) stain^18^. PAS stains polysaccharides (complex sugars like glycogen) such as those found in and around the glomeruli making it a favored stain for delineating them in tissue sections^19^.

The kidney data used in the “HuBMAP - Hacking the Kidney” Kaggle competition comprises 30 whole slide images (WSIs) provided by the BIOmolecular Multimodal Imaging Center (BIOMIC) team at Vanderbilt University (VU) who are also members of HuBMAP’s Tissue Mapping Center at VU (TMC-VU). The tissue blocks were collected through the Cooperative Human Tissue Network^20^ and either fresh frozen (FF) or formalin fixed, paraffin embedded (FFPE)^21^ for preservation. FF tissue is frozen in liquid nitrogen (−190°C) within 30-60 minutes after surgical excision; this type of preservation has been the method of choice for transcriptomics and immunohistochemistry; tissue samples are often embedded in Optimal Cutting Temperature (OCT) media for thin sectioning^22^ or carboxymethylcellulose (CMC) for imaging mass spectrometry^23^. FFPE tissue is the preferred method for clinical pathology samples for histology assessment since the formalin aldehyde cross links proteins to maintain structural integrity of the sample^24^. After preservation, the tissue blocks were sectioned^25^ and imaged using Periodic acid-Schiff (PAS) staining^26^. The slides were scanned with a brightfield scanner, and the resulting images were converted from vendor formats to Tagged Image File Format (TIFF). The images have a spatial resolution of 0.5μm, and the average annotation area was calculated in pixels and μm^2^. On average, the 7,102 glomerulus annotations cover 81,813.5 pixels, or 20,453.4 μm^2^.

Each of the 30 kidney datasets used in the Kaggle competition included a PAS stain whole slide image, anatomical region (AR) masks, and glomeruli segmentation masks. The masks were modified GeoJSON files that captured the polygonal outline of annotations by their pixel coordinates (see samples in **Fig. 1b**), and they were generated from a mix of manually and deep learning (DL) generated annotations. The initial annotations were generated automatically by a segmentation pipeline^27^, then they were inspected and edited by subject matter experts (SMEs)^28^ using QuPath^29^. In addition, information on sample size, location, and rotation within the kidney and pertinent clinical metadata (age, sex, ethnicity, BMI, laterality) was provided (see **Supplementary Table 3**).

For the Kaggle competition, this data was split into three datasets: public train (*n*=15, for training models), public test (*n*=5, for model validation), and private test (*n*=10, for scoring and ranking models). The public datasets were openly available for the competitors to use when designing their models and creating submissions, and the private test set was only available to the Kaggle team and hosts for evaluation of the submissions. After the competition concluded, all data was made available publicly at the HuBMAP Data Portal^15^ as the “HuBMAP ‘Hacking the Kidney’ 2021 Kaggle Competition Dataset - Glomerulus Segmentation on Periodic acid-Schiff Whole Slide Images” collection^30^.

#### Colonic crypts data

Colonic crypts are epithelial invaginations into the connective tissue (stroma) surrounding the colon, or large intestine^31^. Also known as the crypts of Leiberkühn, they contain stem/progenitor cells in their base and are thought to protect these cells from metabolites^32^. They are also the site of absorption and secretion activities within the colon^33^. Normal human colonic crypts have a diameter of 73.5±3.4μm and length of 433±25μm^34^. In addition to stem cells, there are many epithelial subtypes, major subsets include: Paneth (CL:0009009), goblet (CL:1000321), enteroendocrine (CL:0000164), and enterocytes (CL:0002071)^31^. Total number of goblet cells is increasing from the proximal to distal ends of the colon^35^. Enterocytes are absorptive cells which decrease in numbers from the proximal to distal end of the colon and are responsible for absorption of nutrients^35^. Enteroendocrine cells make up a small proportion of the colonic epithelium (<1%) and secrete hormones that control gut physiology^35^.

The colon dataset was provided by the HuBMAP TMC-Stanford team. It consists of two TIFF WSIs and their GeoJSON annotations of colonic crypts. Each image is from a different donor and contains scans of four unique hematoxylin and eosin (H&E) stained coverslips from different regions of the colon (ascending, transverse, descending, and descending sigmoid) for a total of 8 colon H&E images. Hematoxylin and eosin stain nucleic acids deep blue-purple and nonspecific proteins varying degrees of pink, respectively^36^. The two WSIs were annotated by Dr. Teri Longacre using QuPath^29^ and the Manual Annotation of Tissue SOP^37^. The resulting annotations were exported to GeoJSON format and included 395 individual crypt annotations, which on average had an area of 21,331.3 pixels, or 16.1 μm^2^, a considerably smaller average area than that of the glomeruli annotations (81,813.5 pixels, or 20,453.4 μm^2^), see **Supplementary Table 4** for metadata.

#### Spatial location in human body

The HuBMAP Registration User Interface (RUI)^16,38^ was used to capture the three-dimensional size, position, and rotation of all tissue blocks used in this study in close collaboration with subject matter domain experts. The resulting data was used to compute the vertical position of the mass points of all kidney tissue blocks as a proxy of the sequence of tissue sections used here. For the colon, we report the sequence of tissue sections according to the serial extraction sites (ascending colon, transverse colon, descending colon, sigmoid colon).

#### Computation of FTU density

The approximate number of glomerulus annotations in a square millimeter of cortex annotation, henceforth referred to as “FTU density”, was calculated to compare it across cohorts of donors who varied in sex, age, race, and BMI. The 30 glomerulus annotation masks were read into a jupyter notebook from .json format and saved as shapely Polygons^39^. The average area per glomerulus annotation per sample was calculated in pixels, then converted to square microns. The anatomical region masks, which are rough estimates based on quickly-drawn annotations by SMEs, were read into the same jupyter notebook from .json files as shapely polygons, then the total cortex annotation area per sample was calculated by summing the area of all cortex annotations, then converting from pixels to square microns. The approximate FTU density was calculated from these two values and converted to the number of glomerulus annotations per square millimeter.

#### Postprocessing of prediction masks

The 70 prediction masks for all 14 WSI times five algorithm runs were manually examined and FTUs that were overlapping or adjacent were separated via manual addition of a line, see details in Segmentation Mask Analysis section.

### Kidney glomerulus segmentation prior work

For glomerulus segmentation, Sheehan et al. implemented a classifier trained on PAS stain murine renal images through Ilastik^12^. It performed well on their mouse validation set , but when applied to human data it divided glomeruli and generated many false positives. Gallego et al. used transfer learning to fine tune the pre-trained AlexNet CNN with an overlapping sliding window method to segment and classify glomeruli in human WSIs of PAS stained renal tissue. They discovered that the pre-trained model outperformed the model trained from scratch^5^. Govind et al. employed a Butterworth bandpass filter to segment glomeruli from multimodal images (autofluorescence and immunofluorescence marker stain)^40^. Kannan et al. also employed a CNN with an overlapping sliding window operator to segment glomeruli in trichrome-stained images, but they used training data of human origin and watershed segmentation^4^. Methods employing CNNs for the task of glomerulus segmentation seem to be increasingly popular in recent years with highly promising performance^6–8^.

### Colon crypt segmentation prior work

In 2010, Gunduz-Demir et al. approached the task of automatic segmentation of colon glands using an object-graph in conjunction with a decision tree classifier, which obtained a Dice coefficient of 88.91±4.63, an improvement over the pixel-based counterparts at the time^41^. Five years later, Cohen et al. developed a memory-based active contour method that used a random forest classifier that performed pixel level classification with an F-measure of 96.2%^42^. That same year, the Gland Segmentation (GlaS) Challenge Contest was held in conjunction with the Medical Image Computing and Computer-Assisted Intervention (MICCAI 2015) convention^43^. Teams were challenged to present their solutions for automating segmentation of benign and malignant crypts within 165 images from 16 Hematoxylin and Eosin (HE) stained intestinal tissue sections, known as the Warwick-QU dataset. Chen et al. had the winning submission, dubbed “CUMedVision”, which was a novel deep contour-aware fully convolutional neural network (CNN)^44^. Kainz, Pfeiffer, and Urschler submitted the “vision4GlaS” method, a CNN for pixel-wise segmentation and classification paired with a contour based approach to separate pixels into objects, to the GlaS Challenge Contest. Their method ranked 10th in the challenge’s entries^45^. They paired two distinct CNNs (Object-Net for predicting labels and Separator-Net for separating glands) together for pixel-wise classification of the same HE stained images^9^. For this second method, they also preprocessed the RBG images, only inputting the red channel into the model. Banwari et al. took a very computationally efficient approach to colonic crypt segmentation by also isolating the red channel from the GlaS Challenge dataset images and applying intensity based thresholding^11^. Li et al. also used a portion of the GlaS Challenge dataset in 2016 to craft their model, a combination of a window based classification CNN and hand-crafted features with support vector machines (HC-SVM)^46^. Sirinukunwattana, Snead, and Rajpoot used the GlaS challenge dataset in 2015 to develop a random polygons model^47^. In 2018, Tang, Li, and Xu’s Segnet model for crypt segmentation outperformed the contest winner in some portions of the challenge^48^. One of the most recent uses of the GlaS Challenge dataset was by Graham et al. in 2019 for the development of their Minimal Information Loss Dilated Network (MILD-Net) segmentation method which performs simultaneous crypt and lumen segmentation. Their proposed network “counters the loss of information caused by max-pooling by re-introducing the original image at multiple points within the network.” and received higher evaluation metric scores than the winner of the GlaS Challenge or Segnet^49^. Another use of the GlaS Challenge dataset was by Rathore et al. as they tested the efficacy across institutions of their support vector machine (SVM) method for segmenting colonic crypts^50^.

### Competition design

The “HuBMAP - Hacking the Kidney” Kaggle competition teams were tasked with the challenge of detecting glomeruli FTUs in colon data across different tissue preparation pipelines (FF and FFPE). The goal was the implementation of a highly accurate and robust FTU segmentation algorithm.

Two separate types of prizes were offered: Accuracy Prizes and Judges Prizes. The Accuracy Prize awarded $32,000 to the three teams with the highest scores on the Kaggle leaderboard at the conclusion of the competition (1st: $18,000, 2nd: $10,000, 3rd: $4,000). The Judges Prize awarded $28,000 to the teams that advanced science and/or technology (Scientific Prize: $15,000), were the most innovative (Innovation Prize: $10,000), or were the most diverse (Diversity Prize: $3,000) as identified by the panel of judges through a presentation of the teams’ findings and subsequent scoring based on a predetermined rubric^51^. Teams were allowed to enter in multiple categories and had the option of either receiving cash prizes or choosing to have their winnings donated to a charity foundation. Additionally, the use of supplemental publicly available training data was allowed, but organizers were not permitted to participate.

The competition launched on November 16th, 2020 and ran through the final submission date of May 10th, 2021. The data was updated and timeline extended on March 9th, 2021, and the Awards ceremony was held on May 21st, 2021. Submissions were made in the form of Kaggle notebooks with a run-length encoding of the predictions saved in a “submission.csv” file. The notebooks had to run in less than or equal to 9 hours without internet access. See Competition Rules^52^ and Judging Rubric^51^ for more details.

Algorithm performance was evaluated using the mean Dice coefficient (see Metrics). The leaderboard scores were the mean of the Dice coefficients for all ten WSI in the private test set. Any test WSI with predictions missing completely were factored into the mean score as a zero. This metric has been successfully used for previous segmentation task challenges. For example, 922 teams competed in the “Ultrasound Nerve Segmentation” Kaggle competition^53^. The top scoring teams achieved a mean Dice coefficient of 0.73226 and 0.73132 for the private and public leaderboards, respectively. Another competition, entitled “SIIM-ACR Pneumothorax”, engaged 1,475 teams to classify and segment pneumothorax from chest radiographic images, with leaderboard scores topping at 0.8679 and 0.9304 mean Dice coefficients for private and public datasets, respectively^54^. A third competition,”Severstal: Steel Defect Detection” focused on localizing and classifying surface defects on a sheet of steel^55^; it had 2,427 teams competing and achieved mean Dice coefficients of 0.90883 (private leaderboard) and 0.92472 (public leaderboard).

In the “HuBMAP - Hacking the Kidney” Kaggle competition, a total of 1,200 teams competed and the top-5 scoring teams had a mean Dice coefficient of 0.9515 and 0.9512 for the private and public leaderboards, respectively. These are the highest scores for this type of challenge ever achieved.

### Transfer learning

While the Kaggle competition involved developing models for segmenting glomeruli in kidney tissue samples, it is crucial to test the generalization capability of such segmentation models across other organs. To accomplish this goal, we implemented several strategies to train and test the models: 1) The models are trained only on the kidney data and tested on kidney data. 2) The models are trained on kidney data and tested on colon data (without training on any colon data). 3) The models are trained only on colon data and tested on colon data. 4) The models are trained on colon data (using the pretrained models on kidney data for transfer learning) and tested on colon data.

The fourth strategy is called transfer learning in machine learning. It is widely used to improve performance on a dataset by pretraining it on a different but similar dataset. This allows the model to learn more features from the previous dataset and helps improve the generalizability of the overall model. Transfer learning may involve training the entire model or freezing some layers of the model and training the remaining unfrozen layers.

### Algorithms

Teams “Tom,” “Gleb,” and “Whats goin on” won first, second and third place for the accuracy prize respectively. DeepLive.exe and Deepflash2 won the first and second judges prizes respectively. The setup, optimization, and prediction run of all five algorithms are discussed here.

#### Tom

The model uses a single U-Net SeResNext101 architecture with Convolutional Block Attention Module (CBAM)^56^, hypercolumns, and deep supervision. It reads the WSIs as tiled 1024×1024 pixel images and then further resized as 320×320 tiles and sampled using a balanced sampling strategy. The model is trained using a combination of Binary Cross-entropy loss^57^ and Lovász Hinge loss^58^, and the optimizer used is SGD (Stochastic gradient descent)^59^. Training is for 20 epochs, with a learning rate of 10^−4^ to 10^−6^ and batch size of 8 (i.e., training is done using batches of 8 samples per batch).

For the model trained on colon data from scratch or using transfer learning, the training is done for 50-100 epochs and the validation set is increased from 1 slide to 2 slides.

#### Gleb

The model is trained using an ensemble of four 4-fold models namely, Unet-regnety16, Unet-regnetx32, UnetPlusPlus-regnety16^60^, and Unet-regnety16 with scse attention decoder. The model reads tiles of size 1024×1024 sampled from the kidney/colon data. During model training, general data augmentation techniques such as adding gaussian blur and sharpening, adding gaussian noise, applying random brightness or gamma value are used. The models are trained for 50-80 epochs each, with a learning rate of 10^−4^ to 10^−6^, and batch size of 8. The loss function is Dice coefficient loss^61^ and the optimizer used is AdamW^62^.

For the model trained on data from scratch or using transfer learning, the model is trained for 50-100 epochs and the sampling downscale factor is changed from 3 to 2.

#### Whats goin on

Model training uses an ensemble of 2 sets of 5-fold models using the U-Net^63^ architecture (pretrained on Image) with resnet50_32×4d and resnet101_32×4d^64^ as backbones, respectively. Additionally, the a Feature Pyramid Network (FPN)^65^ is added to provide skip connections between upscaling blocks of the decoder, atrous spatial pyramid pooling (ASPP)^66^ is added to enlarge receptive fields, and pixel shuffle^67^ is added instead of transposed convolution to avoid artifacts. The model reads kidney/colon data downsampled by a factor of 2 and tiles of size 1024×1024 are sampled and filtered based on a saturation threshold of 40. General data augmentation techniques are used such as flipping, rotation, scale shifting, deformation, artificial blurring, Hue Saturation Value (HSV) shifting, Contrast Limited Adaptive Histogram Equalization (CLAHE), brightness and contrast shifting, and Piecewise Affine. The models are trained for 50 epochs each, using a one cycle learning rate scheduler with pct_start=0.2, div_factor=1e2, max_lr=1e-4, batch size of 16. The model uses an expansion tile size of 32. The model uses binary cross entropy loss, with gradient norm clipping at 1 and Adam optimizer.

For the model trained on data from scratch or using transfer learning, the batch size is increased to 64 and the expansion tile size is increased to 64.

#### DeepLive.exe

The model architecture used is a simple U-net^68^ with an efficientnet-b^69^ encoder. In addition to the provided training data, the model is trained on additional data from Mendeley^70^ (31 WSIs), Zenodo^71^ (20 WSIs), and the HuBMAP Data Portal^15^ (2 WSIs). The additional data is annotated into two classes: healthy and unhealthy glomeruli. The model employs a dynamic sampling approach whereby it samples tiles of size 512×512 pixels (at a resolution downscale factor of 2) and 768×768 pixels (at a resolution downscale factor of 3). The tiles are sampled from regions having visible glomeruli in them based on annotations, instead of sampling randomly. Model training uses the cross-entropy loss, Adam optimizer, an adaptive learning rate (linearly increased up to 0.001 during the first 500 iterations and then linearly decreased to 0), and a batch size of 32. During training the general data augmentation techniques are used such as brightness and contrast changes, RGB shifting, HSV shifting, color jittering, artificial blurring, CutMix^72^ and MixUp^73^. The model is trained using 5-fold cross validation for at least 10,000 iterations. The key to the model is to reframe the problem as a healthy/unhealthy glomerulus classification problem along with a segmentation problem. This setup enables the model to learn to classify the unhealthy glomeruli as glomeruli and then decide whether the particular instance is healthy enough.

For the model trained on colon data from scratch, on_spot_sampling of 1 and an overlap factor of 2 is used. For the model trained on colon data using transfer learning, on_spot_sampling is set to 1 and an overlap factor of 1 is used. In both cases, no external datasets are used for training.

#### Deepflash2

The model architecture used is a simple U-Net architecture with an efficientnet-b2 encoder (pretrained on ImageNet^74^). Input data is converted and stored as .zarr file format for efficient loading on runtime. The model collectively employs two sampling approaches: 1) Sampling tiles that contain all glomeruli (to ensure that each glomerulus is seen at least once during each epoch of training). 2) Sampling random tiles based on region (cortex, medulla, background) probabilities (to give more weight to the cortex region during training since glomeruli are mainly found in the cortex). The region sampling probabilities were chosen based on expert knowledge and experiments: 0.7 for cortex, 0.2 for medulla, and 0.1 for background. On runtime, the model samples tiles of size 512×512 and uses a resolution downscale factor of 2, 3, and 4 in subsequent runs. During training, general data augmentation techniques are applied such as flipping, blurring, deformation, etc. Model training uses a weighted sum of Dice^75^ and cross-entropy loss^76^ (where both losses have equal weight), Ranger^77^ optimizer (a combination of RAdam^78^ and LookAhead optimizer^79^), a maximum learning rate of 1e-3, and a batch size of 16. The model training is done using a learning rate scheduler whereby the learning rate is scheduled with a cosine annealing^80^ from max_learning_rate / div to max_learning_rate (where div=25). The models are trained and tested using 5-fold cross validation in which each fold is trained on 12 WSIs and validated on 3 WSIs. The best model ensemble for the final score consists of three models trained on different zoom scales (i.e., 2x, 3x, 4x).

For the model trained on the colon data (both with and without transfer learning), the background probability is set to 0.1 and the colon probability is set to 0.9 for sampling, since the colon data lacks the masks for anatomical structures. A weight decay of 10^−5^ was added (for the model trained without transfer learning). For the transfer learning model, saved weights are loaded from the model trained on kidney data at 3x downsampling and the first 13 parameter groups are frozen during training.

### Performance Metrics Terminology

#### Ground Truth

The set of all FTU segmentations in the human annotated dataset using the SOP at (cite) is called ground truth (GT, blue in **Fig. 4**).

**Figure 4.**
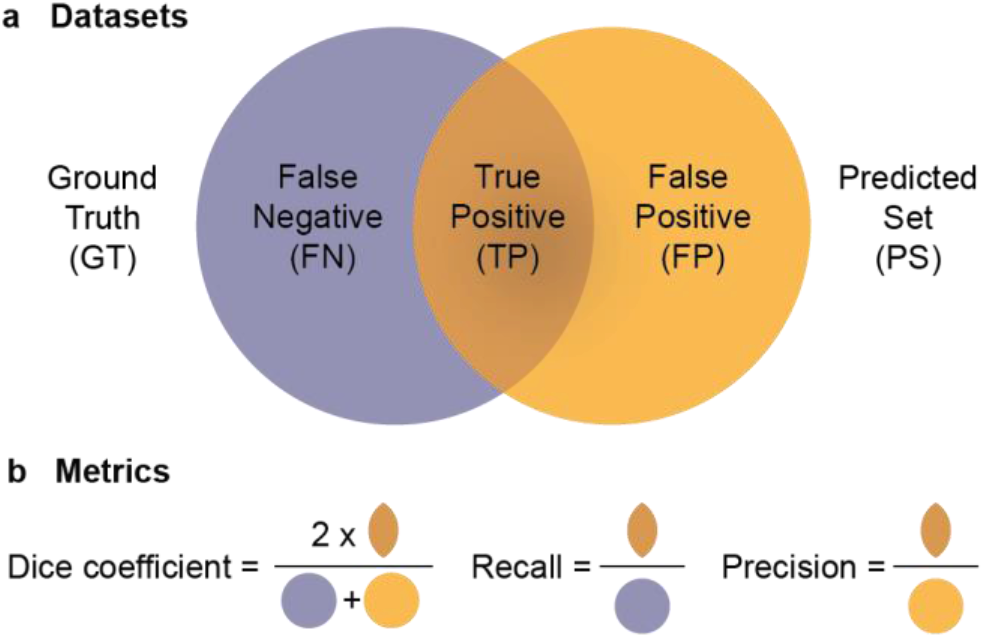
Performance metrics terminology. **a.** Ground truth, predicted set, and false negatives, true positives and false positives datasets. **b.** Dice coefficient, recall, and precision metrics.

#### Predicted Set

The set of all FTU segmentations predicted by an algorithm is called the predicted set (PS, purple in **Fig. 4**).

#### False Negatives, True Positives, and False Positives

Typically, the GT and PS sets overlap creating three sets that are called false negatives (FN, FTUs not predicted by the algorithm), true positives (TP, FTUs in ground truth that are correctly predicted by the algorithm), and false positives (FP, FTUs predicted by the algorithm but not present in the ground truth), see **Fig. 4**).

The sets can be represented via vector-based polylines or pixel masks and different algorithms are used to compare these. Note that the metrics in **Fig. 4** can be applied to pixels that represent an object of interest (e.g., an FTU) or to FTU counts.

### Performance Metrics

**Dice coefficient**, or Sørensen–Dice index^81^, is widely used to compare the pixel-wise agreement between a predicted segmentation and its corresponding ground truth. The formula is given by 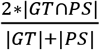, see **Fig. 4**. The Dice coefficient is defined to be 1 when both sets are empty.
**Mean Dice coefficient** is the sum of all Dice coefficients (e.g., one for each image in the test set) divided by the count of all numbers in the collection (e.g., the number of images in the test set).
**Recall**, also referred to as **sensitivity**, measures the proportion of instances that were correctly predicted compared to the sum of false negatives and true positives. It is defined as 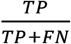, see **Fig. 4**.
**Precision** denotes the proportion of predictions that were correct and it is defined as 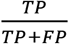, see **Fig. 4**.

Other performance metrics used by related work, see **Supplementary Table 1 and 2**:

#### F-measure/F-score/F_1_-score

The F-measure, also called the F-score or F_1_-score is the harmonic mean of Precision and Recall, defined as 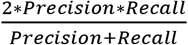.

#### Accuracy

Accuracy is the proportion of of correct predictions as defined by 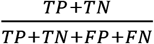

#### Matthews correlation coefficient

The Matthews correlation coefficient is used for binary classifiers to provide a balanced measure of quality^4^. It is defined as 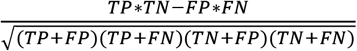.

#### Hausdorff Distance

The Hausdorff distance is a measure used to calculate how similar two objects or images are to one another by calculating the distance between two sets of edge points ^82^.

#### Jaccard index

The Jaccard index, also known as Intersection over Union (IoU), is defined by 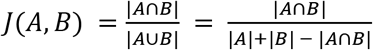, where *A* and *B* are the two objects being compared, e.g., GT and PS in **Fig. 4**. It represents the proportion of area of overlap out of the area of union for the two objects.

### Segmentation Mask Analysis

Ground truth segmentation masks were provided as vector files (one polyline per FTU; many FTUs per WSI). However, algorithm predictions are generated as run-length encodings—one mask for all FTUs in each WSI. Some FTUs are adjacent, effectively merging multiple FTUs into one; this makes it hard to count FTUs or to compute the Dice coefficient but also recall and precision per FTU.

Manually, we added 647 lines to the 70 predicted kidney WSI segmentation masks (232 lines for 50 kidney slides and 415 lines for 20 colon slides) to separate glued together FTUs. We then converted pixel masks for each FTU into one polyline per FTU. Next, we calculated the Dice coefficient for each segmented FTU (glomerulus or crypt) separately; assuming that a Dice coefficient greater than 0.5 indicates that the FTU was correctly predicted, the set of true positives. All FTUs with a Dice coefficient less than 0.5 are false positives (FP), while all ground truth masks with no matching algorithm predictions are false negatives (FN). All results of Dice coefficient, recall, and precision computations are provided in **Supplementary Table 5**.

## Acknowledgements

We are deeply indebted to the generosity of the “HuBMAP - Hacking the Kidney” Kaggle sponsors Google HCLS, Roche, Deloitte, Deerfield Healthcare, Pistoia Alliance, American Chemical Society (CAS), Maven Wave for their contributions of award money for the winning competitors. We thank Andrea de Souza and Deepika Vuppalanch of Eli Lilly and Company for obtaining sponsors, publicising the event, and participating in the awards ceremony. We are also grateful to InnovationDigi/Conference Ventures for launching the competition in conjunction with their conference. We thank Richard Holland, Marcos Novaes, and Addison Howard of the Google team for support throughout the competition. We are also grateful to the Kaggle Scientific prize judges Zorina Galis (NIH), Thomas Fuchs (Paige.ai, Mount Sinai), Matt Nelson (Deerfield Healthcare), Amy Bernard (Allen Institute), Maigan Brusko (University of Florida), Blue Lake (UC San Diego), David Van Valen (CalTech), and Alex Wolf (Cellarity) for sharing their time and expertise while judging the Kaggle submissions. We thank Bruce W. Herr II (Indiana University) for his assistance with collecting and compiling the kidney and colon tissue block data inside the Exploration User Interface. We would also like to thank Bill Shirey (University of Pittsburgh) for his support in registering tissue blocks via the HuBMAP Ingest Portal. This research has been funded in part by the NIH Common Fund through the Office of Strategic Coordination/Office of the NIH Director under award OT2OD026671, NIH awards U54EY032442-01 and U54HG010426-01, National Institute of Diabetes and Digestive and Kidney Diseases (NIDDK) award U54DK120058, and by the NIDDK Kidney Precision Medicine Project grant U2CDK114886. The content is solely the responsibility of the authors and does not necessarily represent the official views of the National Institutes of Health.

## Author contributions

LLG aided the design and implementation of the Kaggle competition. Acted as liaison to Kaggle competitors. Co-wrote the paper. YJ co-designed the Kaggle competition. Ran transferred learning (model training and fine-tuning) for Tom and Gleb algorithms. Implemented metrics for algorithm evaluation and visualizations for result analysis. Co-wrote the paper and developed interactive data visualizations. NS ran algorithms on IU infrastructure to reproduce Kaggle results. Ran transfer learning for Whats goin on and Deepflash2 algorithms. Contributed to performance analysis and comparison of the five algorithms. YJ ran algorithm fine-tuning and training for the Deepflash2 model. Co-lead debugging setup/training issues for machine learning models. Co-wrote the paper. Organized code on GitHub. EMQ wrote up a technical description for participants of the Kaggle competition. Provided glomerulus mask quality control and microscopy expertise. Co-wrote the paper. AB supported spatial registration of kidney and colon data. Compiled tissue metadata. Co-wrote and supported submission of paper. Developed an interactive website. TL manually annotated crypts in colon data.

AMH organized compilation of the colon dataset. YL contributed to compilation of colon samples and images including 3D tissue registration. EDE contributed to construction of the colon dataset. JWH imaged data, performed image processing, created H&E images for colon, and contributed to colon data write up. MPS contributed to the conceptualization and implementation of the transfer learning study using colon.

NHP assisted with the design and implementation of the Kaggle competition and generated kidney FTU segmentation masks. Contributed to kidney data writeup. JMS assisted with the design and implementation of the Kaggle competition and coordinated efforts in generating kidney images and segmentation masks. KB led the design and implementation of the Kaggle competition. Co-lead algorithm comparison and transfer learning. Co-wrote the paper.

## Data availability

Kidney data used in the Kaggle competition has been published via the HuBMAP portal as a collection at https://doi.org/10.35079/hbm925.sgxl.596. The colon data and supplemental tables can be downloaded from https://github.com/cns-iu/ccf-research-kaggle-2021.

## Code availability

Code to re-run the algorithm comparison, compute metrics, and reproduce figures is available at https://github.com/cns-iu/ccf-research-kaggle-2021.

## Supplementary table legends

All tables can be accessed at <u>Supplementary Tables</u>.

**Supplementary table 1.**
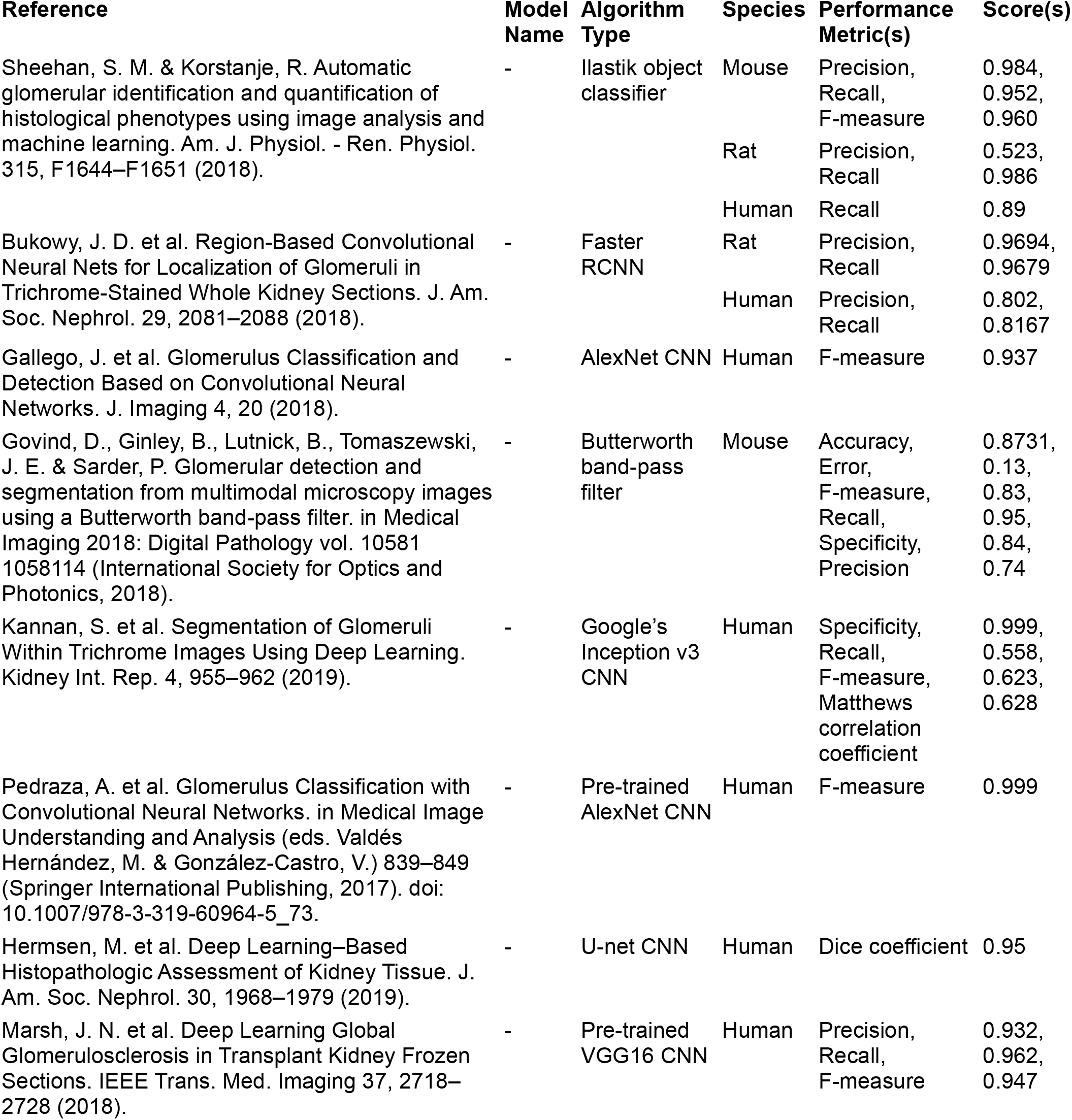
Prior work on renal glomerulus segmentation. This table lists prior work on renal glomerulus segmentation. For each published paper given in the Reference column, we list model name (if applicable), algorithm type, tissue donor species, performance metrics used, and scores achieved.

**Supplementary table 2.**
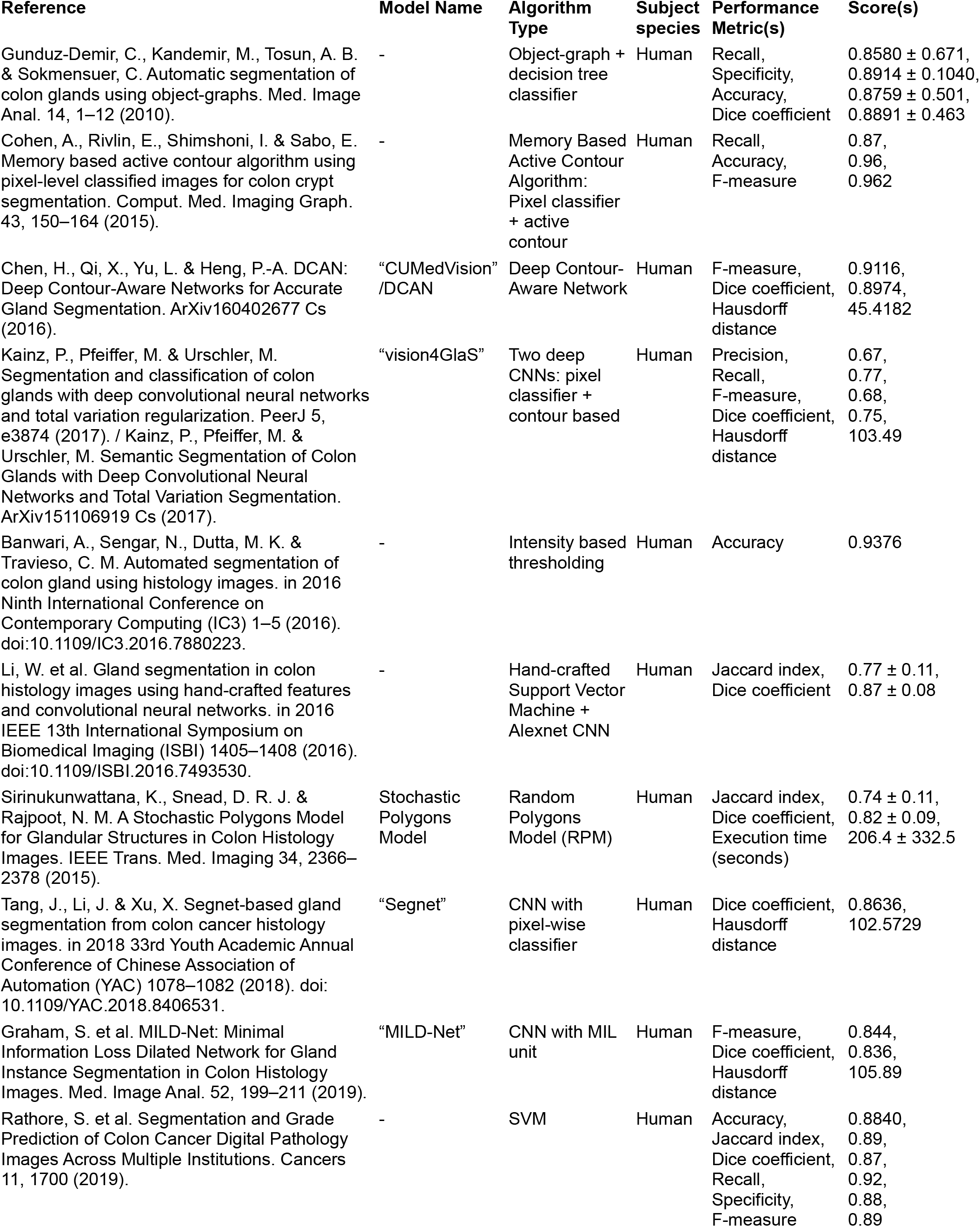
Prior work on colon crypts segmentation. This table lists prior work on colon crypts segmentation. For each published paper given in the Reference column, we list model name (if applicable), algorithm type, tissue donor species, performance metrics used, and scores achieved.

**Supplementary table 3.** Kidney metadata. This table provides metadata for all 30 kidney WSI, one per row. For each WSI, we assigned a running number ID that is also used in **Fig. 1** and **3**. We provide corresponding HuBMAP sample and donor IDs, as well as the Kaggle IDs. We list tissue preservation method (fresh frozen, FF; formalin fixed, paraffin embedded, FFPE), image width and height in pixels, race (White, W; Black or African American, B), sex (male, M; female, F), weight, height, BMI, age, laterality (right kidney, R; left kidney, L), tissue block vertical location (y-position) according to Registration User Interface (RUI) registration. We also computed the glomerulus annotation area in square microns and the approximate number of glomeruli per square millimeter of kidney cortex. Note that there is one patient with 4 tissue blocks in this dataset, 3 patients with 3 blocks, 5 patients who have 2, and the rest only have one. Of the 30 tissue blocks, 25 are from white patients and 5 are from black or African American patients. The dataset is evenly divided between Male/Female sources. All females sampled were white, but male samples were split between white (10) and black or african american (5). None of the samples were associated with a Hispanic or Latino ethnicity. All samples came from adults (minimum age 31 years old). The average weight (85.9kg) lies between the average weights for females (77.47kg) and males (90.63kg) in the United States^83^. The average height (170.35cm) also lies between the average heights for females (161.29cm) and males (175.26cm) in the US^83^. The average BMI of the dataset (29.61kg/m^2^) is between that given as the average for females (29.8kg/m^2^) and males (29.4kg/m^2^) in the US^83^. Only 6 of the samples fell into the “Healthy weight” category (18.5– 24.9), and they originated from two patients. The other 24 samples were either in the “Overweight” (25.0–29.9, 8 samples) or “Obese” (30 and above, 16 samples) categories. There are no noticeable abnormalities when comparing weight and height between the sexes. Average BMI was still above “healthy” (24.9) for each subset when sex was taken into account.

| Slide | HuBMAP ID | Donor ID | Kaggle ID | Tissue preservation method | Image width (pixels) | Image height (pixels) | Race | Sex | Weight (kg) | Height (cm) | BMI | Age | Laterality | Tissue block y-position in 3D reference organ | Number of glomeruli | Average glomerulus area (square micron) | Approximate number of glomeruli per square mm cortex |
| --- | --- | --- | --- | --- | --- | --- | --- | --- | --- | --- | --- | --- | --- | --- | --- | --- | --- |
| 1 | HBM874.RZDW.757 | HBM679.GXQW.326 | 095bf7a1f | FF | 39000 | 38160 | W | F | 71.7 | 160 | 28 | 44 | R | 11.08 | 350 | 24855 | 2.39 |
| 2 | HBM463.JRTB.582 | HBM938.LVRS.434 | aa05346ff | FF | 47340 | 30720 | W | F | 59 | 160 | 23 | 58 | R | 11.69 | 325 | 29153 | 2.06 |
| 3 | HBM636.ZPTS.368 | HBM938.LVRS.434 | b9a3865fc | FFPE | 40429 | 31295 | W | F | 59 | 160 | 23 | 58 | R | 13.70 | 469 | 14081 | 4.22 |
| 4 | **HBM324.ZGZM.874** | **HBM485.HTBW.247** | **e464d2f6c** | **FFPE** | **40816** | **50560** | **W** | **F** | **91.6** | **165.1** | **33.6** | **57** | **R** | **13.93** | **315** | **22774** | **1.96** |
| 5 | **HBM832.FQKR.463** | **HBM485.HTBW.247** | **bacb03928** | **FF** | **22163** | **23968** | **W** | **F** | **91.6** | **165.1** | **33.6** | **57** | **R** | **14.11** | **118** | **14096** | **2.74** |
| 6 | **HBM649.XFQG.775** | **HBM485.HTBW.247** | **ff339c0b2** | **FFPE** | **38912** | **48544** | **W** | **F** | **91.6** | **165.1** | **33.6** | **57** | **R** | **14.28** | **341** | **22522** | **2.34** |
| 7 | HBM958.GHFM.676 | HBM938.LVRS.434 | afa5e8098 | FF | 43780 | 36800 | W | F | 59 | 160 | 23 | 58 | R | 16.35 | 235 | 28262 | 1.63 |
| 8 | **HBM296.RLWW.755** | **HBM485.HTBW.247** | **a14e495cf** | **FFPE** | **32768** | **62688** | **W** | **F** | **91.6** | **165.1** | **33.6** | **57** | **R** | **20.31** | **355** | **20946** | **2.29** |
| 9 | HBM623.RPMC.638 | HBM455.HLHM.985 | 3589adb90 | FFPE | 22165 | 29433 | W | F | 71.3 | 167.6 | 25.4 | 66 | L | 5.91 | 239 | 13868 | 4.93 |
| 10 | HBM276.PGFS.693 | HBM769.HVDR.369 | 2f6ecfcdf | FFPE | 25794 | 31278 | W | F | 93 | 157.4 | 37.5 | 76 | L | 9.58 | 160 | 12455 | 3.27 |
| 11 | HBM849.XMPC.398 | HBM455.HLHM.985 | 26dc41664 | FF | 42360 | 38160 | W | F | 71.3 | 167.6 | 25.4 | 66 | L | 10.81 | 245 | 26441 | 2.57 |
| 12 | HBM362.PTQJ.743 | HBM758.JRSC.348 | d488c759a | FF | 29020 | 46660 | W | F | 81.5 | 158.8 | 32.2 | 66 | L | 11.15 | 175 | 20966 | 2.08 |
| 13 | **HBM673.JJRZ.435** | **HBM633.KPHW.963** | **5274ef79a** | **FF** | **18491** | **22134** | **W** | **F** | **74.6** | **162.6** | **28.2** | **77** | **L** | **12.42** | **51** | **11891** | **1.92** |
| 14 | HBM979.HDZH.896 | HBM769.HVDR.369 | 57512b7f1 | FF | 43160 | 33240 | W | F | 93 | 157.4 | 37.5 | 76 | L | 18.72 | 141 | 24352 | 1.59 |
| 15 | HBM875.QHDJ.259 | HBM547.NCQL.874 | aaa6a05cc | FFPE | 13013 | 18484 | W | F | 87.5 | 162.3 | 33.2 | 73 | L | 99.49 | 99 | 10835 | 4.52 |
| 16 | **HBM344.LLLV.539** | **HBM429.BVWN.357** | **5d8b53a68** | **FF** | **36732** | **22153** | **W** | **M** | **116.9** | **182.9** | **34.9** | **62** | **R** | **9.75** | **320** | **14675** | **3.2** |
| 17 | **HBM627.RSGW.898** | **HBM226.XVDP.877** | **00a67c839** | **FFPE** | **28672** | **30400** | **W** | **M** | **96.6** | **175.3** | **31.4** | **66** | **R** | **14.02** | **97** | **22860** | **1.16** |
| 18 | **HBM227.THVC.544** | **HBM226.XVDP.877** | **0749c6ccc** | **FFPE** | **26624** | **30368** | **W** | **M** | **96.6** | **175.3** | **31.4** | **66** | **R** | **14.18** | **109** | **22929** | **1.55** |
| 19 | HBM783.GJWP.694 | HBM368.WSHR.356 | 0486052bb | FFPE | 34937 | 25784 | W | M | 106.1 | 180.3 | 32.6 | 31 | R | 14.33 | 130 | 18331 | 2.51 |
| 20 | HBM783.GDKK.879 | HBM522.WZBV.379 | 1e2425f28 | FF | 32220 | 26780 | W | M | 131.5 | 193 | 35.3 | 48 | R | 18.05 | 178 | 26214 | 1.89 |
| 21 | HBM833.DBGG.252 | HBM322.KQBK.747 | 2ec3f1bb9 | FFPE | 47723 | 23990 | W | M | 91.2 | 167.6 | 32.5 | 56 | L | 50.70 | 399 | 14134 | 2.85 |
| 22 | HBM389.MBWW.346 | HBM745.MDSR.597 | e79de561c | FF | 27020 | 16180 | B | M | 73 | 166 | 26.5 | 53 | L | 51.24 | 180 | 22025 | 2.6 |
| 23 | **HBM649.DLZF.463** | **HBM322.KQBK.747** | **1eb18739d** | **FF** | **33103** | **20329** | **W** | **M** | **91.2** | **167.6** | **32.5** | **56** | **L** | **54.63** | **157** | **16385** | **2.49** |
| 24 | HBM662.PMPZ.644 | HBM322.KQBK.747 | c68fe75ea | FF | 19780 | 26840 | W | M | 91.2 | 167.6 | 32.5 | 56 | L | 56.86 | 118 | 27664 | 0.92 |
| 25 | HBM879.CDHB.995 | HBM745.MDSR.597 | b2dc8411c | FFPE | 31262 | 14844 | B | M | 73 | 166 | 26.5 | 53 | L | 57.83 | 138 | 11058 | 4.05 |
| 26 | **HBM984.PMZN.942** | **HBM525.JNPV.685** | **9e81e2693** | **FF** | **33100** | **27642** | **B** | **M** | **79.9** | **190.5** | **22** | **58** | **L** | **61.97** | **175** | **25669** | **2.35** |
| 27 | HBM264.XSVF.528 | HBM687.KPKM.763 | 4ef6695ce | FF | 50680 | 39960 | W | M | 91.4 | 181.6 | 27.7 | 56 | L | 134.43 | 439 | 25405 | 1.97 |
| 28 | HBM676.SNVK.793 | HBM687.KPKM.763 | 8242609fa | FFPE | 44066 | 31299 | W | M | 91.4 | 181.6 | 27.7 | 56 | L | 136.11 | 586 | 13800 | 3.55 |
| 29 | HBM636.GVWP.354 | HBM525.JNPV.685 | cb2d976f4 | FFPE | 49548 | 34940 | B | M | 79.9 | 190.5 | 22 | 58 | L | 138.44 | 319 | 19704 | 2.79 |
| 30 | HBM725.PDDC.788 | HBM525.JNPV.685 | 54f2eec69 | FF | 22240 | 30440 | B | M | 79.9 | 190.5 | 22 | 58 | L | 138.91 | 139 | 25465 | 2.63 |

**Supplementary table 4.**
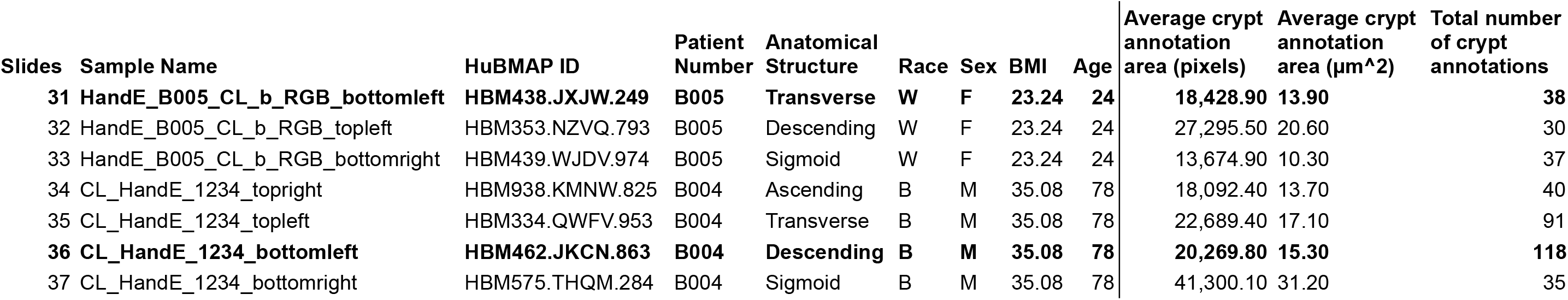
Colon metadata. This table provides metadata for the seven colon WSI. Four of these dataset were sampled from a male donor and three from a female donor. For each WSI, we assigned a running number ID that is also used in **Fig. 1**. We provide corresponding HuBMAP sample and donor IDs, and the Kaggle IDs. We list race (White, W; Black or African American, B), sex (male, M; female, F), BMI, and age. We also computed the average crypt annotation area in square microns.

**Supplementary table 5.** Algorithm performance. This table lists DICE coefficients, false negatives (FN), true positives (TP), and false positives (FP) of winning algorithms for individual WSIs in all three predicted datasets (10 WSI kidney in Kaggle reproduced task, 2 WSI colon in transfer learning task (trained on kidney and colon; tested on colon), 2 WSI colon in from scratch task (trained on colon; tested on colon). Threshold used for calculations is 0.5.

| Kaggle reproduced (kidney) |  |  |  |  |  |  |  |  |  |  |  |  |  |  |  |  |  |  |  |  |  |  |  |
| --- | --- | --- | --- | --- | --- | --- | --- | --- | --- | --- | --- | --- | --- | --- | --- | --- | --- | --- | --- | --- | --- | --- | --- |
| Slide | HuBMAP ID | Ground truth<br>#FTUs | Tom |  |  |  | Gleb |  |  |  | Whats goin on |  |  |  | DeepLive.exe |  |  |  | Deepflash2 |  |  |  | DICE accross 5 models |
|  |  |  | DICE | FN | TP | FP | DICE | FN | TP | FP | DICE | FN | TP | FP | DICE | FN | TP | FP | DICE | FN | TP | FP | DICE |
| 4 | HBM324.ZGZM.874 | 315 | 0.97 | 1 | 314 | 4 | 0.97 | 2 | 313 | 4 | 0.97 | 1 | 314 | 4 | 0.97 | 1 | 314 | 6 | 0.96 | 0 | 315 | 9 | 0.97 |
| 5 | HBM649.DLZF.463 | 157 | 0.95 | 3 | 154 | 3 | 0.95 | 5 | 152 | 5 | 0.95 | 3 | 154 | 6 | 0.95 | 3 | 154 | 5 | 0.95 | 2 | 155 | 8 | 0.95 |
| 6 | HBM296.RLWW.755 | 355 | 0.97 | 2 | 353 | 2 | 0.97 | 4 | 351 | 6 | 0.97 | 3 | 352 | 7 | 0.97 | 1 | 354 | 5 | 0.96 | 1 | 354 | 7 | 0.97 |
| 8 | HBM649.XFQG.775 | 341 | 0.97 | 7 | 334 | 4 | 0.97 | 7 | 334 | 4 | 0.97 | 7 | 334 | 5 | 0.97 | 7 | 334 | 7 | 0.96 | 2 | 339 | 8 | 0.97 |
| 13 | HBM673.JJRZ.435 | 51 | 0.93 | 2 | 49 | 2 | 0.93 | 2 | 49 | 2 | 0.93 | 1 | 50 | 2 | 0.92 | 1 | 50 | 4 | 0.92 | 0 | 51 | 4 | 0.93 |
| 16 | HBM344.LLLV.539 | 320 | 0.93 | 9 | 311 | 14 | 0.93 | 16 | 304 | 31 | 0.93 | 16 | 304 | 18 | 0.93 | 7 | 313 | 23 | 0.93 | 11 | 309 | 21 | 0.93 |
| 17 | HBM627.RSGW.898 | 97 | 0.94 | 2 | 95 | 5 | 0.95 | 1 | 96 | 4 | 0.95 | 2 | 95 | 4 | 0.95 | 2 | 95 | 5 | 0.94 | 1 | 96 | 8 | 0.95 |
| 18 | HBM227.THVC.544 | 109 | 0.96 | 1 | 108 | 2 | 0.96 | 4 | 105 | 3 | 0.96 | 3 | 106 | 2 | 0.96 | 1 | 108 | 3 | 0.96 | 1 | 108 | 3 | 0.96 |
| 23 | HBM832.FQKR.463 | 118 | 0.93 | 4 | 114 | 5 | 0.93 | 4 | 114 | 5 | 0.94 | 2 | 116 | 7 | 0.94 | 1 | 117 | 7 | 0.93 | 5 | 113 | 5 | 0.93 |
| 26 | HBM984.PMZN.942 | 175 | 0.96 | 5 | 170 | 2 | 0.94 | 10 | 165 | 5 | 0.95 | 9 | 166 | 3 | 0.95 | 6 | 169 | 7 | 0.95 | 4 | 171 | 8 | 0.95 |
| Transfer learning (trained on kidney & colon, tested on colon) |  |  |  |  |  |  |  |  |  |  |  |  |  |  |  |  |  |  |  |  |  |  |  |
| Slide | HuBMAP ID | Ground truth<br>#FTUs | Tom |  |  |  | Gleb |  |  |  | Whats goin on |  |  |  | DeepLive.exe |  |  |  | Deepflash2 |  |  |  | DICE accross 5 models |
|  |  |  | DICE | FN | TP | FP | DICE | FN | TP | FP | DICE | FN | TP | FP | DICE | FN | TP | FP | DICE | FN | TP | FP | DICE |
| 31 | HBM438.JXJW.249 | 36 | 0.83 | 2 | 34 | 12 | 0.76 | 10 | 26 | 8 | 0.75 | 2 | 34 | 16 | 0.82 | 2 | 34 | 10 | 0.65 | 11 | 25 | 29 | 0.76 |
| 36 | HBM462.JKCN.863 | 124 | 0.93 | 7 | 117 | 5 | 0.91 | 10 | 114 | 2 | 0.89 | 5 | 119 | 11 | 0.93 | 6 | 118 | 9 | 0.78 | 25 | 99 | 46 | 0.89 |
| From scratch (trained on colon, tested on colon) |  |  |  |  |  |  |  |  |  |  |  |  |  |  |  |  |  |  |  |  |  |  |  |
| Slide | HuBMAP ID | Ground truth<br>#FTUs | Tom |  |  |  | Gleb |  |  |  | Whats goin on |  |  |  | DeepLive.exe |  |  |  | Deepflash2 |  |  |  | DICE accross 5 models |
|  |  |  | DICE | FN | TP | FP | DICE | FN | TP | FP | DICE | FN | TP | FP | DICE | FN | TP | FP | DICE | FN | TP | FP | DICE |
| 31 | HBM438.JXJW.249 | 36 | 0.82 | 1 | 35 | 15 | 0.75 | 10 | 26 | 7 | 0.71 | 3 | 33 | 18 | 0.85 | 1 | 35 | 12 | 0.75 | 4 | 32 | 21 | 0.78 |
| 36 | HBM462.JKCN.863 | 124 | 0.93 | 6 | 118 | 10 | 0.90 | 15 | 109 | 6 | 0.88 | 3 | 121 | 11 | 0.94 | 6 | 118 | 3 | 0.86 | 9 | 115 | 11 | 0.90 |

## References

1. Snyder, M. P. et al. The human body at cellular resolution: the NIH Human Biomolecular Atlas Program. Nature 574, 187–192 (2019).

2. de Bono, B., Grenon, P., Baldock, R. & Hunter, P. Functional tissue units and their primary tissue motifs in multi-scale physiology. J. Biomed. Semant. 4, 22–22 (2013).

3. Hayman, J. M., Jr., Martin, J. W., Jr. & Miller, M. Renal Function and the Number of Glomeruli in the Human Kidney. Arch. Intern. Med. 64, 69–83 (1939).

4. Kannan, S. et al. Segmentation of Glomeruli Within Trichrome Images Using Deep Learning. Kidney Int. Rep. 4, 955–962 (2019).

5. Gallego, J. et al. Glomerulus Classification and Detection Based on Convolutional Neural Networks. J. Imaging 4, 20 (2018).

6. Hermsen, M. et al. Deep Learning–Based Histopathologic Assessment of Kidney Tissue. J. Am. Soc. Nephrol. 30, 1968–1979 (2019).

7. Pedraza, A. et al. Glomerulus Classification with Convolutional Neural Networks. in Medical Image Understanding and Analysis (eds. Valdés Hernández, M. & González-Castro, V.) 839–849 (Springer International Publishing, 2017). doi:10.1007/978-3-319-60964-5_73.

8. Marsh, J. N. et al. Deep Learning Global Glomerulosclerosis in Transplant Kidney Frozen Sections. IEEE Trans. Med. Imaging 37, 2718–2728 (2018).

9. Kainz, P., Pfeiffer, M. & Urschler, M. Segmentation and classification of colon glands with deep convolutional neural networks and total variation regularization. PeerJ 5, e3874 (2017).

10. Chen, H. et al. DCAN: Deep contour-aware networks for object instance segmentation from histology images. Med. Image Anal. 36, 135–146 (2017).

11. Banwari, A., Sengar, N., Dutta, M. K. & Travieso, C. M. Automated segmentation of colon gland using histology images. in 2016 Ninth International Conference on Contemporary Computing (IC3) 1–5 (2016). doi:10.1109/IC3.2016.7880223.

12. Sheehan, S. M. & Korstanje, R. Automatic glomerular identification and quantification of histological phenotypes using image analysis and machine learning. Am. J. Physiol. - Ren. Physiol. 315, F1644–F1651 (2018).

13. HuBMAP - Hacking the Kidney. https://kaggle.com/c/hubmap-kidney-segmentation.

14. Börner, K. et al. Anatomical Structures, Cell Types, and Biomarkers Tables Plus 3D Reference Organs in Support of a Human Reference Atlas. bioRxiv 2021.05.31.446440 (2021) doi:10.1101/2021.05.31.446440.

15. HuBMAP Data Portal. https://portal.hubmapconsortium.org/.

16. HuBMAP CCF Registration User Interface (CCF-RUI). https://hubmapconsortium.github.io/ccf-ui/rui/.

17. Mounier-Vehier, C. et al. Cortical thickness: An early morphological marker of atherosclerotic renal disease. Kidney Int. 61, 591–598 (2002).

18. Vaughan, M. R. & Quaggin, S. E. How Do Mesangial and Endothelial Cells Form the Glomerular Tuft? J. Am. Soc. Nephrol. 19, 24–33 (2008).

19. Agarwal, S. K., Sethi, S. & Dinda, A. K. Basics of kidney biopsy: A nephrologist’s perspective. Indian J. Nephrol. 23, 243 (2013).

20. Brewer, M. Collection and Post-Surgical Excision of Human Kidney Tissue through the Cooperative Human Tissue Network. (2019) doi:10.17504/protocols.io.7gehjte.

21. Allen, J. et al. Freezing Fresh Tissue. protocols.io https://www.protocols.io/view/freezing-fresh-tissue-6wghfbw (2019).

22. Robbe, P. et al. Clinical whole-genome sequencing from routine formalin-fixed, paraffin-embedded specimens: pilot study for the 100,000 Genomes Project. Genet. Med. 20, 1196–1205 (2018).

23. Stoeckli, M., Staab, D. & Schweitzer, A. Compound and metabolite distribution measured by MALDI mass spectrometric imaging in whole-body tissue sections. Int. J. Mass Spectrom. 260, 195–202 (2007).

24. Bass, B. P., Engel, K. B., Greytak, S. R. & Moore, H. M. A Review of Preanalytical Factors Affecting Molecular, Protein, and Morphological Analysis of Formalin-Fixed, Paraffin-Embedded (FFPE) Tissue: How Well Do You Know Your FFPE Specimen? Arch. Pathol. Lab. Med. 138, 1520–1530 (2014).

25. Anderson, D. et al. Cryostat Sectioning of Tissues for 3D Multimodal Molecular Imaging. protocols.io https://www.protocols.io/view/cryostat-sectioning-of-tissues-for-3d-multimodal-m-7ethjen (2019).

26. Allen, J. et al. Initial Rapid Pathology Assessment of Kidney Tissue. protocols.io https://www.protocols.io/view/initial-rapid-pathology-assessment-of-kidney-tissu-9dph25n (2020).

27. Patterson, N. H. et al. Autofluorescence microscopy as a label-free tool for renal histology and glomerular segmentation. 2021.07.16.452703 https://www.biorxiv.org/content/10.1101/2021.07.16.452703v1 (2021) doi:10.1101/2021.07.16.452703.

28. Greenwald, N. F. et al. Whole-cell segmentation of tissue images with human-level performance using large-scale data annotation and deep learning. 2021.03.01.431313 https://www.biorxiv.org/content/10.1101/2021.03.01.431313v2 (2021) doi:10.1101/2021.03.01.431313.

29. Bankhead, P. et al. QuPath: Open source software for digital pathology image analysis. Sci. Rep. 7, 16878 (2017).

30. Allen, J. L. et al. HuBMAP ‘Hacking the Kidney’ 2020-2021 Kaggle Competition Dataset - Glomerulus Segmentation on Periodic Acid-Schiff Whole Slide Images. (2021) doi:10.35079/HBM925.SGXL.596.

31. van der Flier, L. G. & Clevers, H. Stem Cells, Self-Renewal, and Differentiation in the Intestinal Epithelium. Annu. Rev. Physiol. 71, 241–260 (2009).

32. Kaiko, G. E. et al. The Colonic Crypt Protects Stem Cells from Microbiota-Derived Metabolites. Cell 165, 1708–1720 (2016).

33. Geibel, J. Secretion and Absorption by Colonic Crypts. Annu. Rev. Physiol. 67, 471–490 (2005).

34. Halm, D. R. & Halm, S. T. Secretagogue response of goblet cells and columnar cells in human colonic crypts1. Am. J. Physiol.-Cell Physiol. 278, C212–C233 (2000).

35. Barker, N., Wetering, M. van de & Clevers, H. The intestinal stem cell. Genes Dev. 22, 1856–1864 (2008).

36. Fischer, A. H., Jacobson, K. A., Rose, J. & Zeller, R. Hematoxylin and Eosin Staining of Tissue and Cell Sections. Cold Spring Harb. Protoc. 2008, pdb.prot4986 (2008).

37. Scherschel, L. & Ju, Y. SOP: Manual Annotation of Tissue. (2021) doi:10.5281/zenodo.5565028.

38. Bueckle, A., Buehling, K., Shih, P. C. & Börner, K. Comparing Completion Time, Accuracy, and Satisfaction in Virtual Reality vs. Desktop Implementation of the Common Coordinate Framework Registration User Interface (CCF RUI). ArXiv210212030 Cs (2021).

39. The Shapely User Manual — Shapely 1.7.1 documentation. https://shapely.readthedocs.io/en/stable/manual.html#polygons.

40. Govind, D., Ginley, B., Lutnick, B., Tomaszewski, J. E. & Sarder, P. Glomerular detection and segmentation from multimodal microscopy images using a Butterworth band-pass filter. in Medical Imaging 2018: Digital Pathology vol. 10581 1058114 (International Society for Optics and Photonics, 2018).

41. Gunduz-Demir, C., Kandemir, M., Tosun, A. B. & Sokmensuer, C. Automatic segmentation of colon glands using object-graphs. Med. Image Anal. 14, 1–12 (2010).

42. Cohen, A., Rivlin, E., Shimshoni, I. & Sabo, E. Memory based active contour algorithm using pixel-level classified images for colon crypt segmentation. Comput. Med. Imaging Graph. 43, 150–164 (2015).

43. Sirinukunwattana, K. et al. Gland Segmentation in Colon Histology Images: The GlaS Challenge Contest. ArXiv160300275 Cs (2016).

44. Chen, H., Qi, X., Yu, L. & Heng, P.-A. DCAN: Deep Contour-Aware Networks for Accurate Gland Segmentation. ArXiv160402677 Cs (2016).

45. Kainz, P., Pfeiffer, M. & Urschler, M. Segmentation and classification of colon glands with deep convolutional neural networks and total variation regularization. PeerJ 5, e3874–e3874 (2017).

46. Li, W. et al. Gland segmentation in colon histology images using hand-crafted features and convolutional neural networks. in 2016 IEEE 13th International Symposium on Biomedical Imaging (ISBI) 1405–1408 (2016). doi:10.1109/ISBI.2016.7493530.

47. Sirinukunwattana, K., Snead, D. R. J. & Rajpoot, N. M. A Stochastic Polygons Model for Glandular Structures in Colon Histology Images. IEEE Trans. Med. Imaging 34, 2366–2378 (2015).

48. Tang, J., Li, J. & Xu, X. Segnet-based gland segmentation from colon cancer histology images. in 2018 33rd Youth Academic Annual Conference of Chinese Association of Automation (YAC) 1078–1082 (2018). doi:10.1109/YAC.2018.8406531.

49. Graham, S. et al. MILD-Net: Minimal Information Loss Dilated Network for Gland Instance Segmentation in Colon Histology Images. Med. Image Anal. 52, 199–211 (2019).

50. Rathore, S. et al. Segmentation and Grade Prediction of Colon Cancer Digital Pathology Images Across Multiple Institutions. Cancers 11, 1700 (2019).

51. HuBMAP - Hacking the Kidney Judging Rubric. https://www.kaggle.com/c/hubmap-kidney-segmentation/overview/judges-prize.

52. HuBMAP - Hacking the Kidney Competition Rules. https://www.kaggle.com/c/hubmap-kidney-segmentation/rules.

53. Ultrasound Nerve Segmentation. https://kaggle.com/c/ultrasound-nerve-segmentation.

54. SIIM-ACR Pneumothorax Segmentation. https://kaggle.com/c/siim-acr-pneumothorax-segmentation.

55. Severstal: Steel Defect Detection. https://kaggle.com/c/severstal-steel-defect-detection.

56. Woo, S., Park, J., Lee, J.-Y. & Kweon, I. S. CBAM: Convolutional Block Attention Module. ArXiv180706521 Cs (2018).

57. Ruby, U. & Yendapalli, V. Binary cross entropy with deep learning technique for Image classification. Int. J. Adv. Trends Comput. Sci. Eng. 9, (2020).

58. Yu, J. & Blaschko, M. The Lovász Hinge: A Convex Surrogate for Submodular Losses. 26.

59. Gardner, W. A. Learning characteristics of stochastic-gradient-descent algorithms: A general study, analysis, and critique. Signal Process. 6, 113–133 (1984).

60. Zhou, Z., Siddiquee, M. M. R., Tajbakhsh, N. & Liang, J. UNet++: Redesigning Skip Connections to Exploit Multiscale Features in Image Segmentation. IEEE Trans. Med. Imaging 39, 1856–1867 (2020).

61. Wang, L., Wang, C., Sun, Z. & Chen, S. An Improved Dice Loss for Pneumothorax Segmentation by Mining the Information of Negative Areas. IEEE Access 8, 167939–167949 (2020).

62. Loshchilov, I. & Hutter, F. Decoupled Weight Decay Regularization. in International Conference on Learning Representations (2019).

63. Ronneberger, O., Fischer, P. & Brox, T. U-Net: Convolutional Networks for Biomedical Image Segmentation. in Medical Image Computing and Computer-Assisted Intervention – MICCAI 2015 (eds. Navab, N., Hornegger, J., Wells, W. M. & Frangi, A. F.) 234–241 (Springer International Publishing, 2015). doi:10.1007/978-3-319-24574-4_28.

64. Xie, S., Girshick, R., Dollár, P., Tu, Z. & He, K. Aggregated Residual Transformations for Deep Neural Networks. in 2017 IEEE Conference on Computer Vision and Pattern Recognition (CVPR) 5987–5995 (2017). doi:10.1109/CVPR.2017.634.

65. Lin, T.-Y. et al. Feature Pyramid Networks for Object Detection. in 2017 IEEE Conference on Computer Vision and Pattern Recognition (CVPR) 936–944 (2017). doi:10.1109/CVPR.2017.106.

66. Chen, L.-C., Papandreou, G., Kokkinos, I., Murphy, K. & Yuille, A. L. DeepLab: Semantic Image Segmentation with Deep Convolutional Nets, Atrous Convolution, and Fully Connected CRFs. IEEE Trans. Pattern Anal. Mach. Intell. 40, 834–848 (2018).

67. Shi, W. et al. Real-Time Single Image and Video Super-Resolution Using an Efficient Sub-Pixel Convolutional Neural Network. in 1874–1883 (2016).

68. Ronneberger, O., Fischer, P. & Brox, T. U-Net: Convolutional Networks for Biomedical Image Segmentation. in Medical Image Computing and Computer-Assisted Intervention – MICCAI 2015 (eds. Navab, N., Hornegger, J., Wells, W. M. & Frangi, A. F.) 234–241 (Springer International Publishing, 2015). doi:10.1007/978-3-319-24574-4_28.

69. Tan, M. & Le, Q. V. EfficientNet: Rethinking Model Scaling for Convolutional Neural Networks. ArXiv190511946 Cs Stat (2020).

70. Bueno, G., Gonzalez-Lopez, L., García-Rojo, M. & Laurinavicius, A. Data for glomeruli characterization in histopathological images. 3, (2020).

71. Swiderska-Chadaj, Z., Gallego, J. & Gertych, A. Kidney glomeruli-ROIs extracted from histological slides stained with HE or PAS. (2020) doi:10.5281/zenodo.4299694.

72. Yun, S. et al. CutMix: Regularization Strategy to Train Strong Classifiers With Localizable Features. in 6023–6032 (2019).

73. Zhang, H., Cisse, M., Dauphin, Y. N. & Lopez-Paz, D. mixup: Beyond Empirical Risk Minimization. ArXiv171009412 Cs Stat (2018).

74. ImageNet. https://image-net.org/.

75. Sudre, C. H., Li, W., Vercauteren, T., Ourselin, S. & Jorge Cardoso, M. Generalised Dice Overlap as a Deep Learning Loss Function for Highly Unbalanced Segmentations. in Deep Learning in Medical Image Analysis and Multimodal Learning for Clinical Decision Support (eds. Cardoso, M. J. et al.) 240–248 (Springer International Publishing, 2017). doi:10.1007/978-3-319-67558-9_28.

76. Cao, J. et al. Softmax Cross Entropy Loss with Unbiased Decision Boundary for Image Classification. in 2018 Chinese Automation Congress (CAC) 2028–2032 (2018). doi:10.1109/CAC.2018.8623242.

77. Wright, L. & Demeure, N. Ranger21: a synergistic deep learning optimizer. ArXiv210613731 Cs (2021).

78. Liu, L. et al. On the Variance of the Adaptive Learning Rate and Beyond. ArXiv190803265 Cs Stat (2020).

79. Zhang, M. R., Lucas, J., Hinton, G. & Ba, J. Lookahead Optimizer: K Steps Forward, 1 Step Back. in Proceedings of the 33rd International Conference on Neural Information Processing Systems (Curran Associates Inc., 2019).

80. Loshchilov, I. & Hutter, F. SGDR: Stochastic Gradient Descent with Warm Restarts. ArXiv160803983 Cs Math (2017).

81. Carass, A. et al. Evaluating White Matter Lesion Segmentations with Refined Sørensen-Dice Analysis. Sci. Rep. 10, (2020).

82. Dong-Gyu Sim, Oh-Kyu Kwon, & Rae-Hong Park. Object matching algorithms using robust Hausdorff distance measures. IEEE Trans. Image Process. 8, 425–429 (1999).

83. Fryar, C., Carroll, M., Gu, Q., Afful, J. & Ogden, C. Anthropometric reference data for children and adults: United States, 2015–2018. 44 https://www.cdc.gov/nchs/data/series/sr_03/sr03-046-508.pdf (2021).

